# Homology Modelling, In Silico Prediction And Characterization Of Cytochrome c oxidase In *Cyprinus carpio* And *Tubifex tubifex* And Molecular Docking Studies Between The Modelled Protein And Three Commonly Used Surfactants Sodium Dodecyl Sulphate, Cetylpyridinium Chloride And Sodium Laureth Sulphate

**DOI:** 10.1101/2021.07.09.451643

**Authors:** Ritwick Bhattacharya, Ismail Daoud, Arnab Chatterjee, Soumendranath Chatterjee, Nimai Chandra Saha

## Abstract

The purpose of this work is to evaluate the homology modeling, in silico prediction, and characterisation of Cytochrome c oxidase from *Cyprinus carpio* and *Tubifex tubifex*, as well as molecular docking experiments between the modelled protein and three frequently used surfactants. Using the template crystal structure of bovine heart Cytochrome c oxidase, homology modeling of Cytochrome c oxidase (Subunit 2) of *Cyprinus carpio* (Accession # P24985) and Cytochrome c oxidase (Subunit 1) of *Tubifex tubifex* (Accession # Q7YAA6) was conducted. The model structure was improved further with 3^D^refine, and the final 3D structure was verified with PROCHEK and ERRATA. The physiochemical, as well as the stereochemical parameters of the modelled protein, were evaluated using various tools like ExPASy’s ProtParam, Hydropathy Analysis and EMBOSS pepwheel. The projected model was then docked with toxic ligands, Sodium dodecyl sulfate (SDS), Cetylpyridinium chloride (CPC), and Sodium laureth sulfate (SLES), whose 3D structures were obtained from the Uniprot database. CPC interacted best with Cytochrome c oxidase subunit 2 of *Cyprinus carpio* and Cytochrome c oxidase subunit 1 of *Tubifex tubifex*, according to our findings. Furthermore, in the case of all surfactants, hydrophobic interactions with the active site amino acid residues of the modelled protein were observed to be more common than hydrogen bonds and salt bridges. Molecular simulation studies exhibited that the surfactants alter the structural flexibility of the predicted proteins. Hence it may be inferred that the surfactants might alter the structure and dynamics of Cytochrome c oxidase of both worm and fish.

## Introduction

Surfactants basically used in personal care products exert detrimental effects on different target as well as non-target organisms which further may also affect humans by biomagnification (Ebele *et al.* 2017). They are perilous to macromolecules and alter their efficient functioning in the biological system by annexing with them (Ivanković and Hrenović 2010). Several studies have documented the toxic effects of surfactants in aquatic organisms (Lechuga *et al.* 2016, Freitas *et al.* 2019, Freitas, Silvestro, Coppola, *et al.* 2020, Freitas, Silvestro, Pagano, *et al.* 2020, Hering *et al.* 2020, Mustapha and Bawa-Allah 2020). The surfactants are entirely categorized into anionic, nonionic, cationic and zwitterionic surfactants (Jackson *et al.* 2016). The most bountiful class of surfactants that are prodigiously utilized in household detergents and industrial cleaning products are anionic surfactants (Ivanković and Hrenović 2010). The most commercially used anionic surfactants especially utilized for household products, cosmetics, and laundry purposes are soidum dodecyl sulphate (SDS) and sodium laureth sulphate (SLES) (Bondi *et al.* 2015, Barra Caracciolo *et al.* 2017). It has been accounted for that the survivability of fishes and microorganisms are antagonistically influenced by anionic surfactants (Chaturvedi and Kumar 2010). On the other hand cationic surfactants are molecules with a long, hydrophobic chain connected to the positive nitrogen atom (Puchta 1984). These are more poisonous contrasted with anionic surfactants (Jardak *et al.* 2016). This group of surfactants is commonly utilized in sundry sectors such as textiles, emulsifiers, wetting agents, disinfectants, and cosmetics (Puchta 1984, Jardak *et al.* 2016). Predicated on the previous studies, it is observed that quaternary ammonium compounds are the most abundant and widely utilized class of cationic surfactant (Zhang *et al.* 2015). Cetylpyridinium chloride (CPC) is indeed a quaternary ammonium cationic component that is deliberately used in mouthwashes for the elimination of dental plaques and periodontitis. Besides, it also eliminates reactive dyes, phenols, and different organic solutes from waste products or treated effluents (Costa *et al.* 2013).

Cytochrome c oxidase (CcO) is the terminal oxidase of the mitochondrial electron transport chain that catalyzes the reduction of molecular oxygen to water (Wikström *et al.* 2018). Any toxicant induced dysfunction or modifications in Cytochrome c oxidase, facilitates oxidative stress by inducing mitochondrial apoptosis (Srinivasan and Avadhani 2012). The protein structure prediction methods include developing a three-dimensional protein structure from its monomer (amine acid). Relatively few protein structures are known in the protein database and the number of sequences has risen considerably faster than the number of protein structure available owing to advancements in DNA sequencing. Homology protein modeling is a systematic approach in the study of protein structure. It gives an insight into the underlying characteristics of proteins and is used to predict the three-dimensional structure (3D). Moreover molecular docking are essential method to identify the effect of hazardous ligands or chemicals by recognizing patterns of inter-molecular interactions with the desired proteins (Dhandare *et al.* 2020).

Several pieces of research were conducted regarding the induction of stress in aquatic organisms upon addition of surfactant (Bhattacharya *et al.* 2019a, 2019b, 2021). Moreover, several researches were also conducted regarding the dysfunction of Cytochrome c oxidase upon addition of several toxicant (Leavesley *et al.* 2008). However, evidence regarding the In silico assessment of the effects of surfactants (SDS, SLES, and CPC) to Cytochrome c oxidase on aquatic organism is meager.

We predicted and analyzed Cytochrome c oxidase from both *Cyprinus carpio* and *Tubifex tubifex* as potential receptor proteins in this work. Based on qualitative and quantitative characteristics, the predicted protein was deemed to be satisfactory and was docked with three commonly used surfactants SDS, CPC, and SLES. Moreover the study was elaborated to note the pattern of interactions between SDS, CPC, SLES and the protein Cytochrome c oxidase, which may serve as a potential implement to assess the effects of the surfactants on stress related proteins in aquatic organisms.

## Materials and methods

### 1. Sequence Retrieval, Modeling and Prediction of target protein

The FASTA sequence of Cytochrome c oxidase (Subunit 2) of *Cyprinus carpio* (Accession # P24985) and Cytochrome c oxidase (Subunit 1) of *Tubifex tubifex* (Accession # Q7YAA6) were retrieved from Uniprot (https://www.uniprot.org/). For homology modeling the 3d structure of the target protein is constructed using SWISS MODEL. It is a fully automated protein structure homology-modelling server, accessible via the Expasy server (Bienert *et al.* 2017, Waterhouse *et al.* 2018). The PDB templates used for the construction of target protein was 3abm.1.B (Cytochrome c oxidase subunit 2 of bovine heart, resolution 1.95 Å) having sequence identity: 71.81% with the modelled protein Cytochrome c oxidase (Subunit 2) of *Cyprinus carpio* and 3abm.1.A (Cytochrome c oxidase subunit 2 of bovine heart, resolution 1.95 Å) having sequence identity 76.23% with the modelled protein Cytochrome c oxidase (Subunit 1) of *Tubifex tubifex.*

### 2. Refinement, Evaluation, and Validation of Protein Model

The predicted protein structures are refined using 3Drefine. It is an online interactive server enabling computationally efficient protein structure refinement with web-based statistics and graphical analysis capabilities. For effective protein structure refinement, the refinement technique employs repeated optimization of the hydrogen bonding network along with atomic-level energy reduction on the optimized model utilizing a combined physics and knowledge-based force fields (Bhattacharya and Cheng 2013). The structures are further evaluated and validated using PROCHECK by constructing Ramachandran plot (Laskowski *et al.* 1993) and ERRAT score (Colovos and Yeates 1993). The Ramachandran plot visualizes the dihedral angles phi and psi of amino acid residues in protein structure and delineates the potential allowed and disallowed configurations of protein structure, whereas the ERRAT interface performs on the statistics of non-bonded atomic interactions and atom allocation (Colovos and Yeates 1993, Laskowski *et al.* 1993).

### 3. Physicochemical Parameter Analysis

ExPASy’s ProtParam method is used to determine different calculated physiochemical parameters of predicted protein such as number of amino acids, molecular weight, theoretical pI, aliphatic index (AI), and grand average of hydropathicity (GRAVY) (Gasteiger *et al.* 1999). Hydropathy Analysis is utilized to determine hydropathy and amphipathicity of the predicted protein, and EMBOSS Pepwheel is used to emphasize amphipathicity and other characteristics of residues around a helix by displaying them in a wheel diagram, which is available in The Transporter Classification Database (Saier *et al.* 2016). GOR IV server is used for prediction of secondary structures of the modelled proteins.

### 4. Retreival of Toxic Ligands

3d conformers of three commonly used surfactants, sodium dodecyl sulphate (SDS), cetylpyridinium chloride (CPC) and sodium laureth sulphate (SLES) were obtained from the PubChem Compound Database (https://pubchem.ncbi.nlm.nih.gov/) in structure data file (sdf) and canonical Simplified Molecular Input Line Entry Specification (SMILES) format.

### 5. Protein-Ligand Docking Study

Prior to docking docking, the predicted protein file was loaded into Biovia Discovery studio for preparation steps. The steps included the addition of polar hydrogen, Kollman charges as well as removal of heteroatoms. Removal of water is not necessary as the presence of water is essential to ensure a relay between the compound and the active site and thus create networks of hydrogen bonds (Klebe 2006). On the other hand water molecules in the cavities of proteins can sometimes be a fundamental element as some algorithms are able to simulate the presence of water molecules in the cavities of proteins (Marechal 2007).The binding affinity of 3 surfactants with target protein is predicted using CB-Dock (Cavity-detection guided Blind Docking) which is designed to perform blind docking at predicted sites, instead of the entire surface of a protein. It implements Autodoc Vina (1.12) for performing docking analysis (Liu *et al.* 2020). The docking pose of the ligand protein complex was selected based on the lowest vina score and RMSD value of less than 2 Angstrom (Liu *et al.* 2020). Finally, the bond type and interaction between active site residues of protein and ligand was obtained by using Protein-Ligand Interaction Profiler (PLIP) which aids in easy and fast identification of intermolecular interactions between biological macromolecules and their ligands (Salentin *et al.* 2015, Adasme *et al.* 2021).

### 6. Molecular Dynamics Simulation studies

The simulations of molecular dynamics (MD) were conducted with the CABS-flex 2.0 server to evaluate the structural flexibilities of the proteins alone and compare it to the protein ligand docking complex (Kurcinski *et al.* 2019). Based on the MD trajectory and default variables the root-mean-square fluctuations (RMSF) were determined. In recent years, several researchers relied on this server for protein ligand simulation studies (Arora *et al.* 2020, Vardhan and Sahoo 2020, Shah *et al.* 2021).

## Results and Discussions

### 1. Structure prediction and model evaluation of target protein

**Fig. 1** shows modelled predicted structure of the target protein, as determined by SWISS MODEL (Waterhouse *et al.* 2018). Ramachandran plot shows the percentage of residues in favourable regions, additionally allowed regions, generously allowed regions, and disallowed regions. The Ramachandran plot obtained through PROCHEK server (Laskowski *et al.* 1993) revealed that our predicted protein model of *Cyprinus carpio* and *Tubifex tubifex* had 90.1% and 94.2% residues respectively, lying in the most allowed region which qualifies it as a good model as according to PROCHEK (a good quality model is expected to have >90% residues in the most favourable region). Moreover, no residues were present in the disallowed region **(Fig 2)**. The overall quality factor and compatibility of the model with amino acid residues were evaluated using ERRAT (Colovos and Yeates 1993) which was calculated as 90.2778 and 94.9622 respectively. In general, the modelled protein is considered good if the quality factor is >50 (Khusro *et al.* 2020).

**Fig 1:**
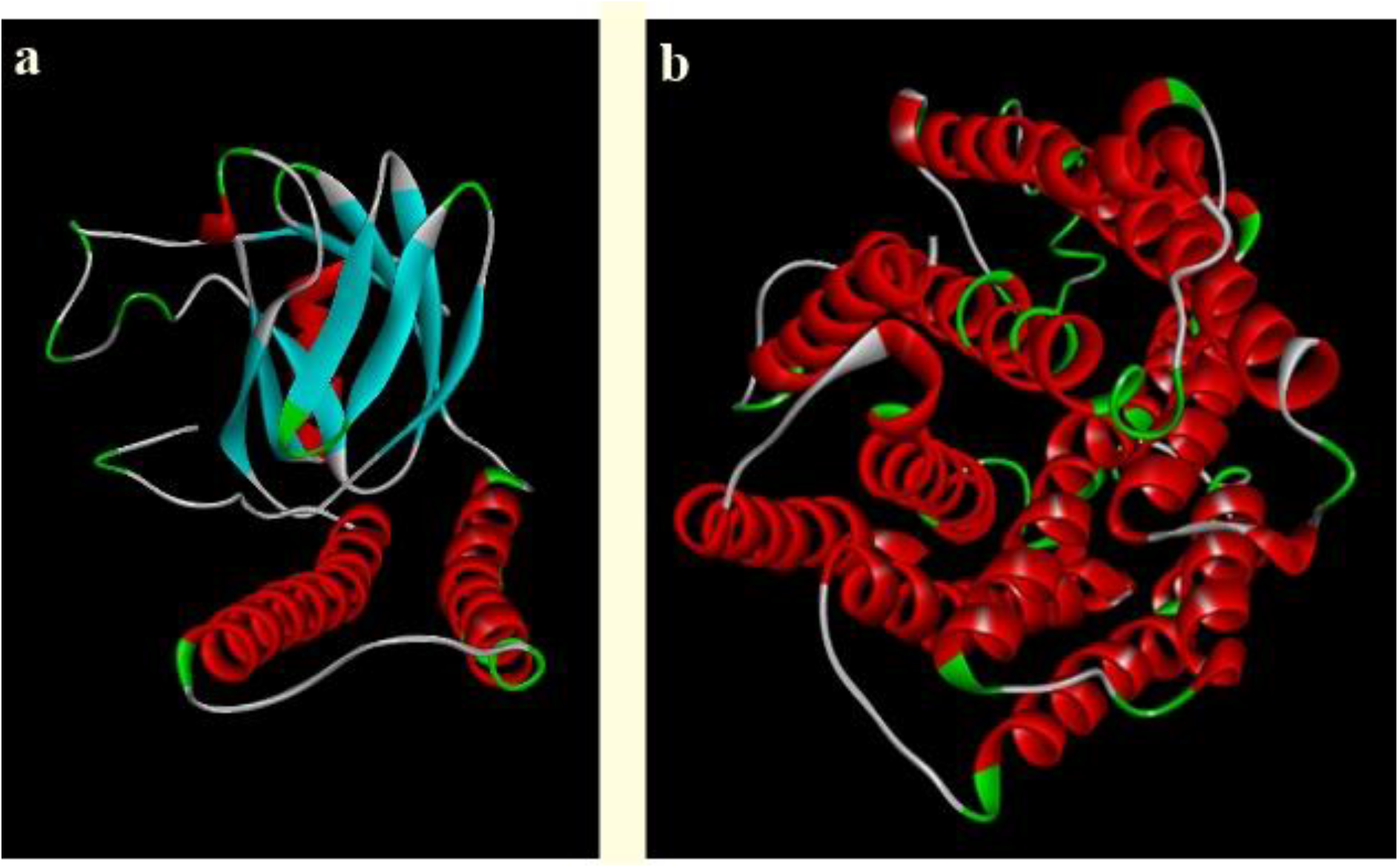
Predicted structure of : **(a):** Cytochrome c oxidase subunit 2 of *Cyprinus carpio* and **(b):**Cytochrome c oxidase subunit 1 (*Tubifex tubifex*).

**Fig 2:**
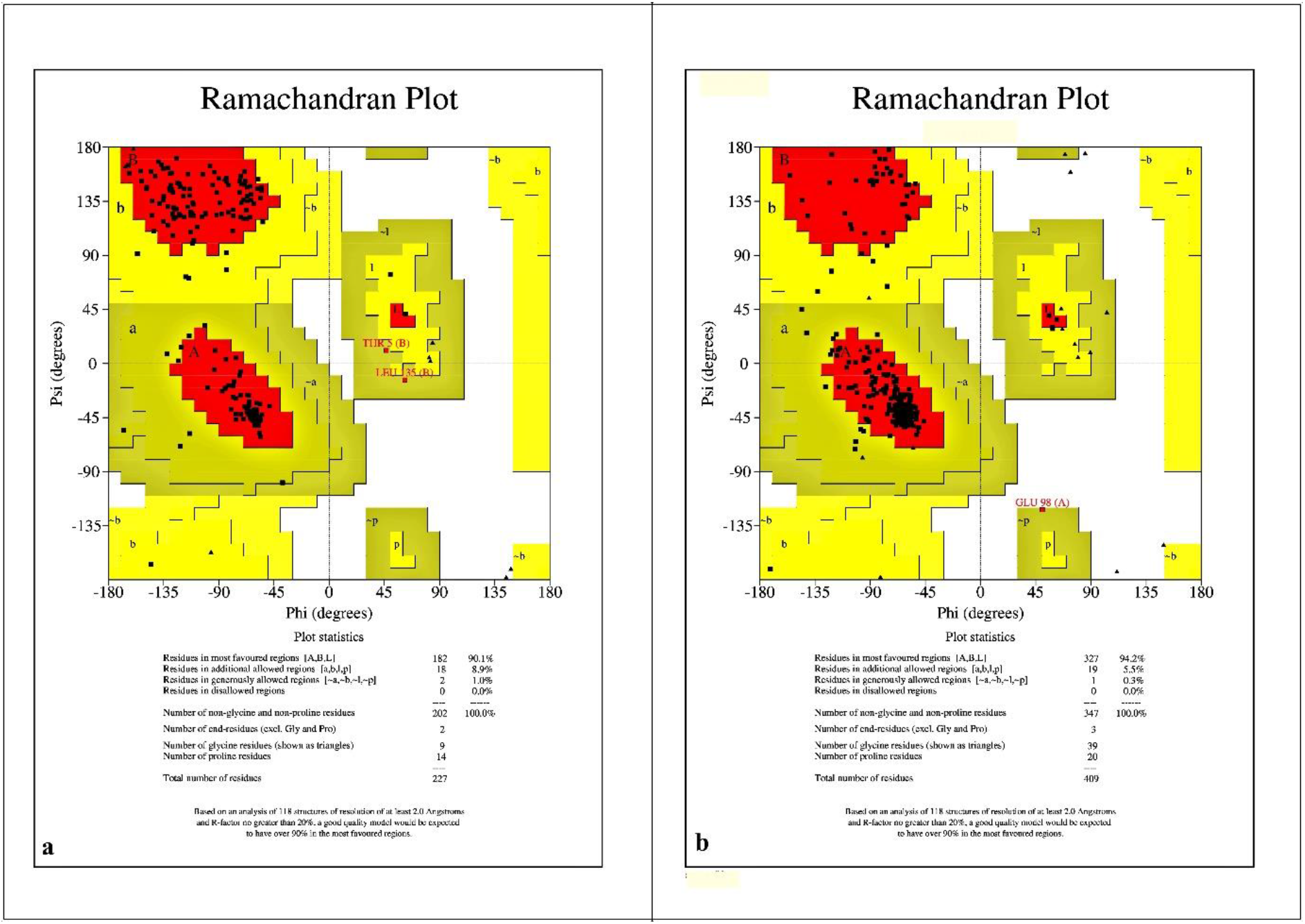
Ramachandran plot of: **(a):** Cytochrome c oxidase subunit 2 of *Cyprinus carpio* and **(b):** Cytochrome c oxidase subunit 1 of *Tubifex tubifex*.

### 2. Physiochemical properties of the modelled proteins

Physiochemical parameters are the preliminary attributes of proteins that exemplify their distinctiveness. (Khusro *et al.* 2020) Results for physiochemical parameters of predicted protein model of *Cyprinus carpio* and *Tubifex tubifex* are shown in **Table 1.** The pH where the surface of a protein is charged but the net charge of the protein is zero is known as the isoelectric point (pI). If the theoretical pI value is more than 7, the macromolecule has an alkaline nature whereas the predicted pI < 7 shows the acidity of the molecule (Shaw *et al.* 2001). In this context, theoretical pI of Cytochrome c oxidase subunit 2 of *Cyprinus carpio* and Cytochrome c oxidase subunit 1 (*Tubifex tubifex*) were determined acidic and alkaline in nature respectively. The aliphatic index (AI) is a measure used to assess protein stability. AI can be determined by the relative volume occupied by the aliphatic side chains of amino acids in a protein. A high aliphatic index value of more than 50 indicates that the protein is temperature stable. In general, the aliphatic index is closely attributed to the protein or peptide’s thermostability (Kaur *et al.* 2020). In this study, the modelled protein Cytochrome c oxidase subunit 2 of *Cyprinus carpio* and Cytochrome c oxidase subunit 1 of *Tubifex tubifex* exhibited the aliphatic index value of 110.61 and 113.31, which indicates that the predicted proteins are highly thermostable. The hydrophobicity value of a peptide is expressed by the grand average of hydropathicity index (GRAVY), which calculates the total of the hydropathy values of all the amino acids divided by the sequence length. Positive GRAVY values imply hydrophobic properties, whereas negative ones suggest hydrophilic properties (Chang and Yang 2013). Based on the GRAVY values, Cytochrome c oxidase subunit 2 of *Cyprinus carpio* and Cytochrome c oxidase subunit 1 of *Tubifex tubifex* are hydrophobic in nature.

**Table 1:** Some Physiochemical parameters of modelled proteins

| Modelled proteins | Physiochemical parameters |  |  |  |
| --- | --- | --- | --- | --- |
|  | Number of amino acids | Theoretical pI | Aliphatic index | GRAVY |
| Cytochrome c oxidase subunit 2<br>(*Cyprinus carpio*) | 230 | 4.79 | 110.61 | 0.327 |
| Cytochrome c oxidase subunit 1<br>(*Tubifex tubifex*) | 408 | 7.44 | 113.31 | 0.825 |

### 3. Primary structure analysis of modelled proteins

The primary structure of peptides reflects the amino acid makeup (Breda *et al.* 2007). The amino acid composition of the modelled proteins Cytochrome c oxidase subunit 2 of *Cyprinus carpio* and Cytochrome c oxidase subunit 1 of *Tubifex tubifex* is shown in **Fig. 3.** In this scenario, Leucine is the most abundant amino acid in the modelled proteins of both Cytochrome c oxidase subunit 2 of *Cyprinus carpio* (with ≈12%) and Cytochrome c oxidase subunit 1 of *Tubifex tubifex* (with ≈14%). Furthermore, the proportion of Glycine in Cytochrome c oxidase subunit 1 of *Tubifex tubifex* (with ≈10%) is greater than Cytochrome c oxidase subunit 2 of *Cyprinus carpio* (with ≈4%) (Khusro *et al.* 2020).

**Fig 3:**
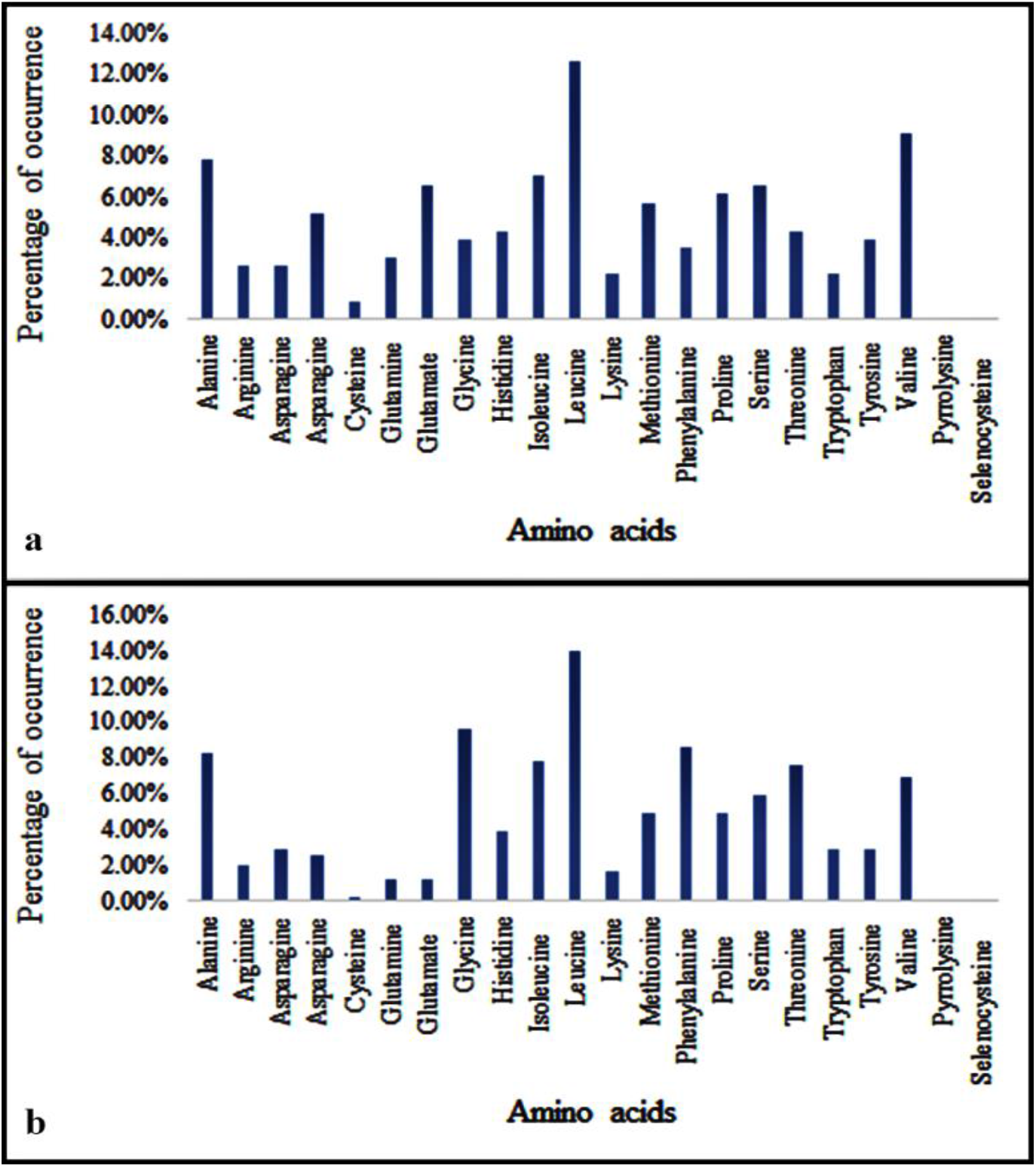
Amino acid composition (%) of: **(a):** Cytochrome c oxidase subunit 2 of *Cyprinus carpio* and **(b):** Cytochrome c oxidase subunit 1 of *Tubifex tubifex*.

### 4. Secondary structure analysis of peptide

The secondary structure of both the modelled proteins of Cytochrome c oxidase subunit 2 of *Cyprinus carpio* and Cytochrome c oxidase subunit 1 of *Tubifex tubifex* consists principally of an alpha helix, an extended strand segment, β-turn, and a random coil **(Fig 4)**. However, the sequence of dominance of the secondary structure in Cytochrome c oxidase subunit 2 of *Cyprinus carpio* is Random coil > Alpha helix > Extended strand > β-turn, but in Cytochrome c oxidase subunit 1 of *Tubifex tubifex* the order of dominance is Alpha helix > Random coil > Extended strand > β-turn. Furthermore, the proportion of β-turn is greater in Cytochrome c oxidase subunit 1 of *Tubifex tubifex* than in Cytochrome c oxidase subunit 2 of *Cyprinus carpio*, indicating that Cytochrome c oxidase subunit 1 of *Tubifex tubifex* has a slightly poorer structural stability than Cytochrome c oxidase subunit 2 of *Cyprinus carpio* as presence of β-turn in protein secondary structure negative impacts the stabiluty of the protein (Khusro *et al.* 2020)

**Fig 4:**
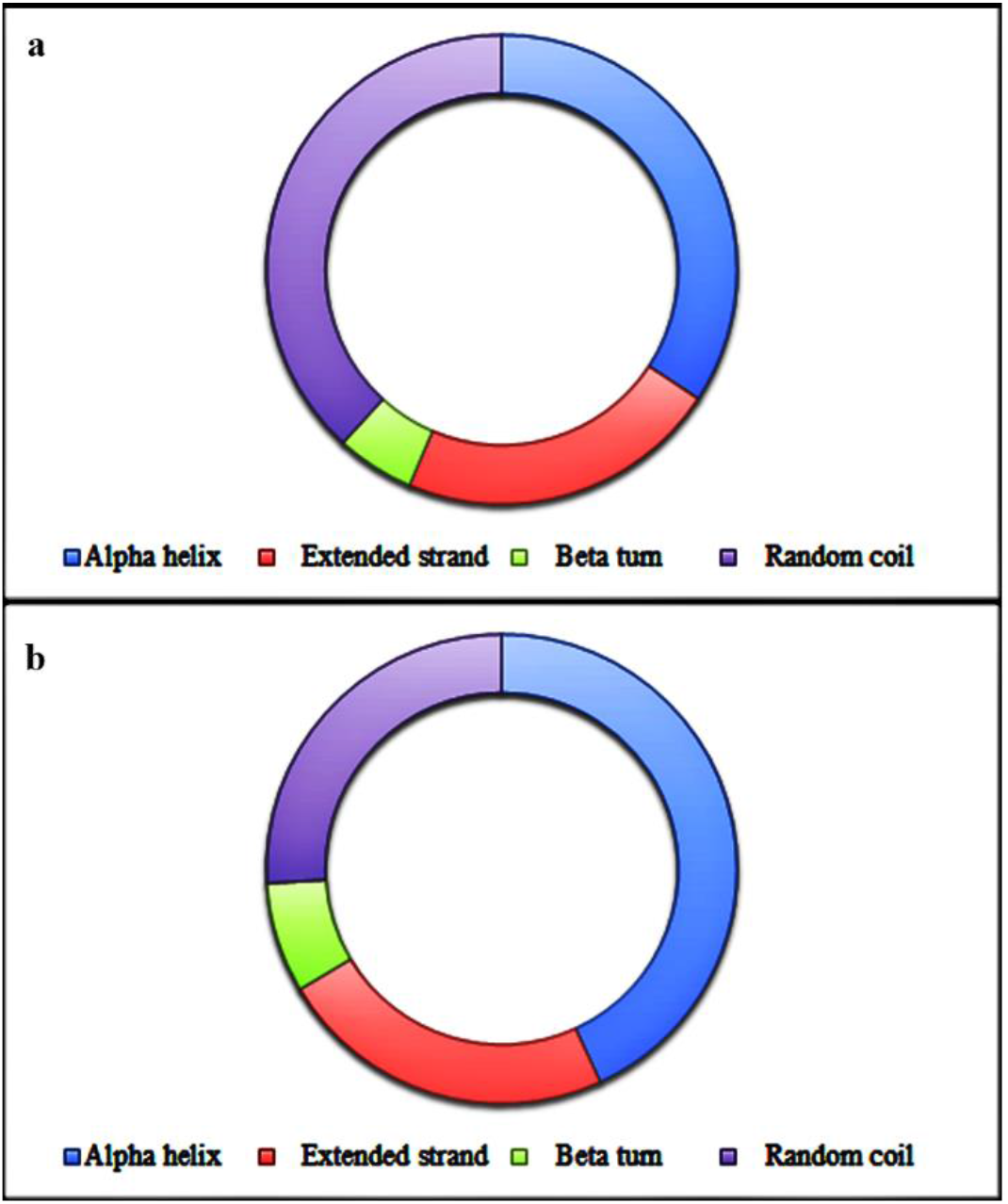
Prevalance of Alpha helix, extended strand region, β-turn, and random coil in **(a):** Cytochrome c oxidase subunit 2 of *Cyprinus carpio*and **(b):** Cytochrome c oxidase subunit 1 of *Tubifex tubifex*

### 5. Functional analysis of modelled protein

Hydropathicity and membrane-spanning properties of Cytochrome c oxidase subunit 2 of *Cyprinus carpio* and Cytochrome c oxidase subunit 1 of *Tubifex tubifex* are illustrated in Fig 5, which showed that Cytochrome c oxidase subunit 2 of *Cyprinus carpio* has two transmembrane regions while Cytochrome c oxidase subunit 1 of *Tubifex tubifex* has ten. This suggests that the frequency of transmembrane segments is greater in Cytochrome c oxidase subunit 1 of *Tubifex tubifex*. Furthermore, the helical wheel diagram depicts the protein sequence in a helical format. It may be used to highlight amphipathicity and other characteristics of residues surrounding a helix (Schiffer and Edmundson 1967). The helical wheel prediction of modelled proteins Cytochrome c oxidase subunit 2 of *Cyprinus carpio* and Cytochrome c oxidase subunit 1 of *Tubifex tubifex* is shown in **Fig. 6,** revealing the allocation of hydrophilic, alipathic, and positively charged residues of Cytochrome c oxidase subunit 2 of *Cyprinus carpio* and Cytochrome c oxidase subunit 1 of *Tubifex tubifex*.

**Fig 5:**
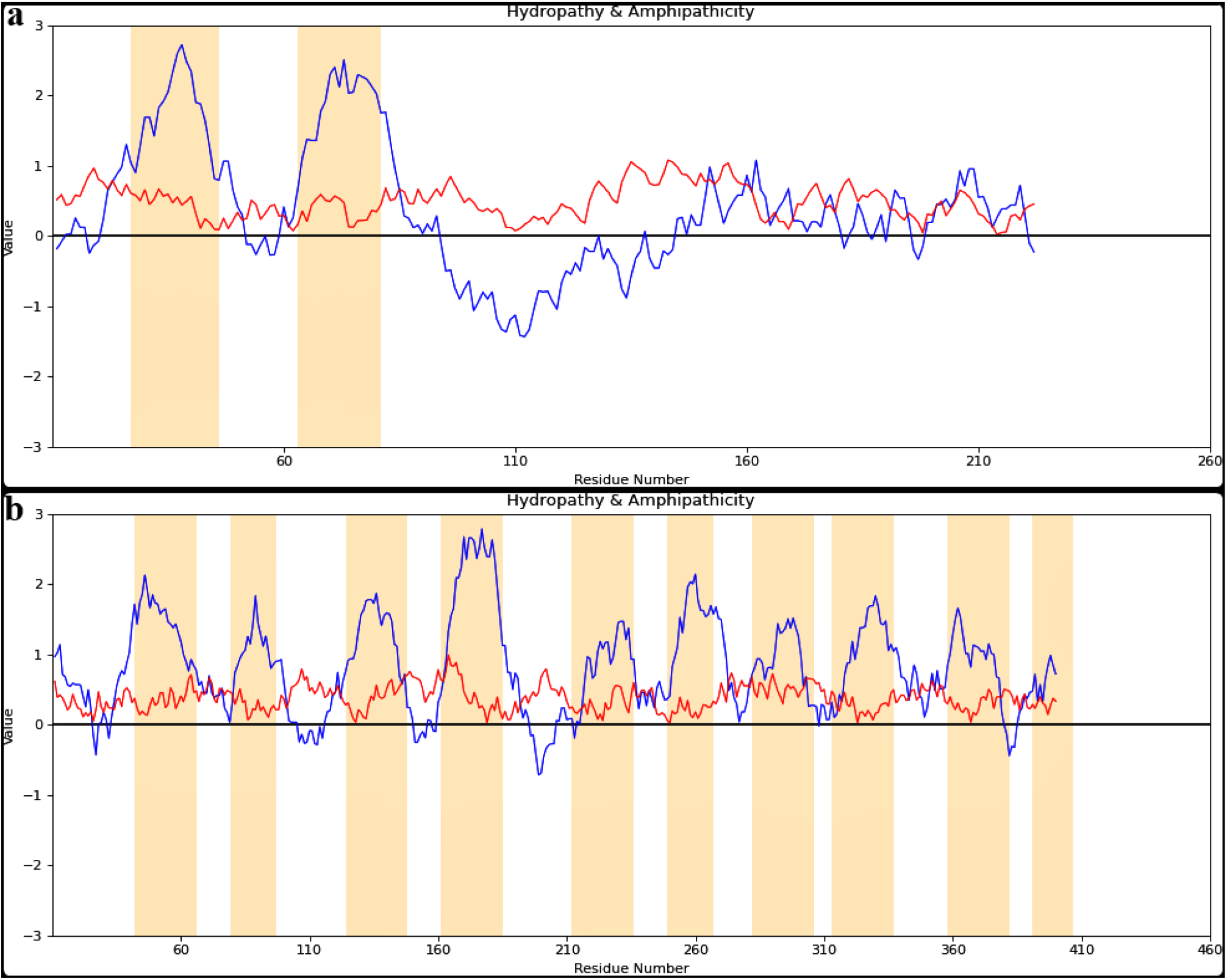
Hydropathicity, amphipathicity and transmembrane segments of **(a):** Cytochrome c oxidase subunit 2 of *Cyprinus carpio* and **(b):** Cytochrome c oxidase subunit 1 of *Tubifex tubifex*. Blue lines denote Hydropathy, Red lines denote Amphipathicity and Orange bars mark transmembrane segments.

**Fig 6:**
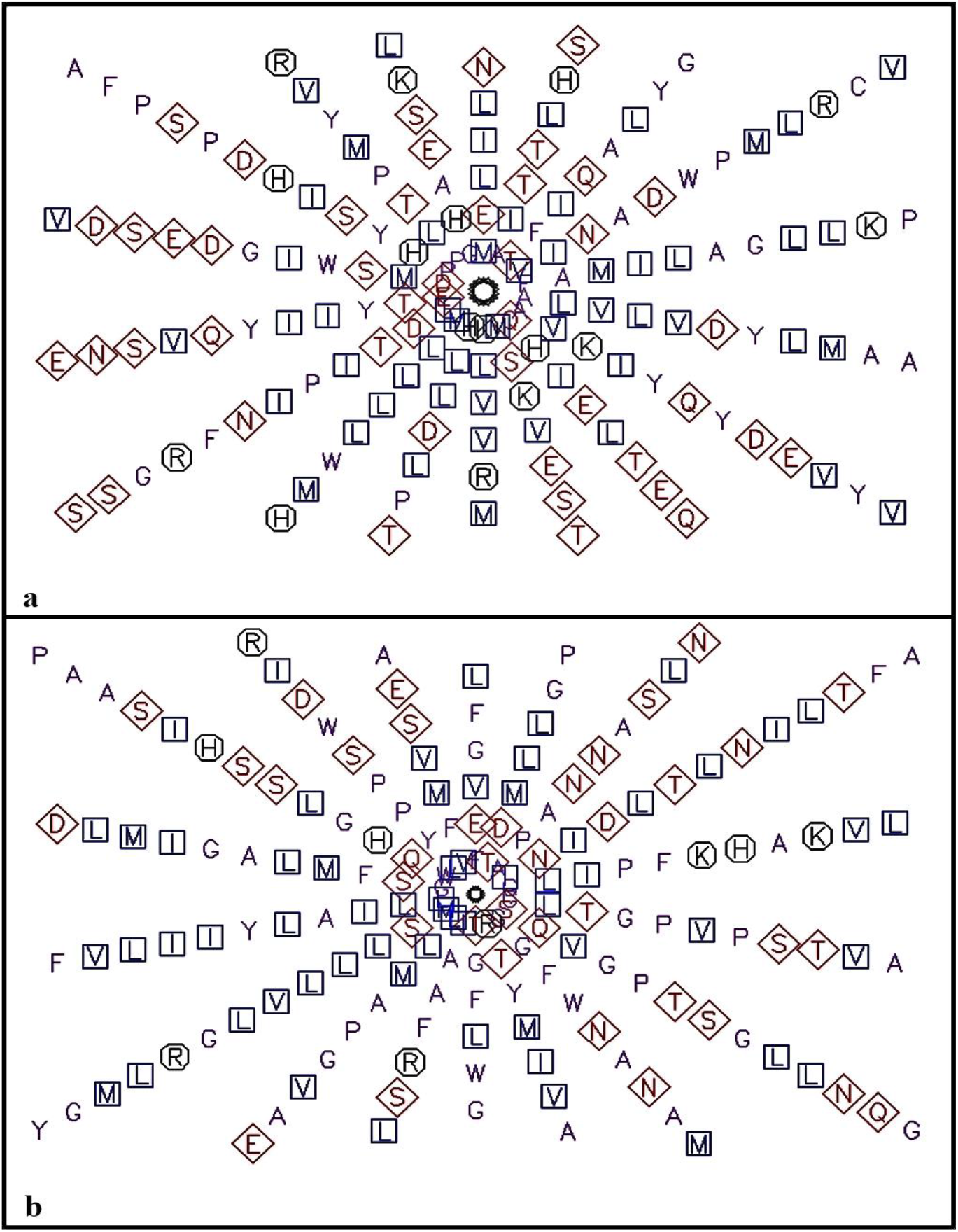
The helical wheel prediction of **(a):** Cytochrome c oxidase subunit 2 of *Cyprinus carpio*and **(b):** Cytochrome c oxidase subunit 1 of *Tubifex tubifex*. Aliphatic residues are marked with squares, hydrophilic residues are marked with diamonds, and positively charged residues with octagons.

### 6. Protein-Ligand Docking

The 3D structure of the surfactant protein interaction in both fish and worm are provided in **Fig. 7.** The docking pose of the ligand protein complex was selected based on the lowest vina score and Root mean square deviation (RMSD) value of less than 2 Å (Liu *et al.* 2020) The root-mean square deviation (RMSD) was used to determine how different the docking orientation achieved is from the equivalent co-crystallized posture of the identical ligand molecule.. Basically three different RMSD classifications are used for docking solutions: (a) good solution when RMSD ≤ 2.0 Å (b) acceptable solutions when RMSD is between 2.0 and 3.0 Å, and (c) bad solutions when RMSD ≥ 3.0 Å (Ramírez and Caballero 2018).

**Fig 7:**
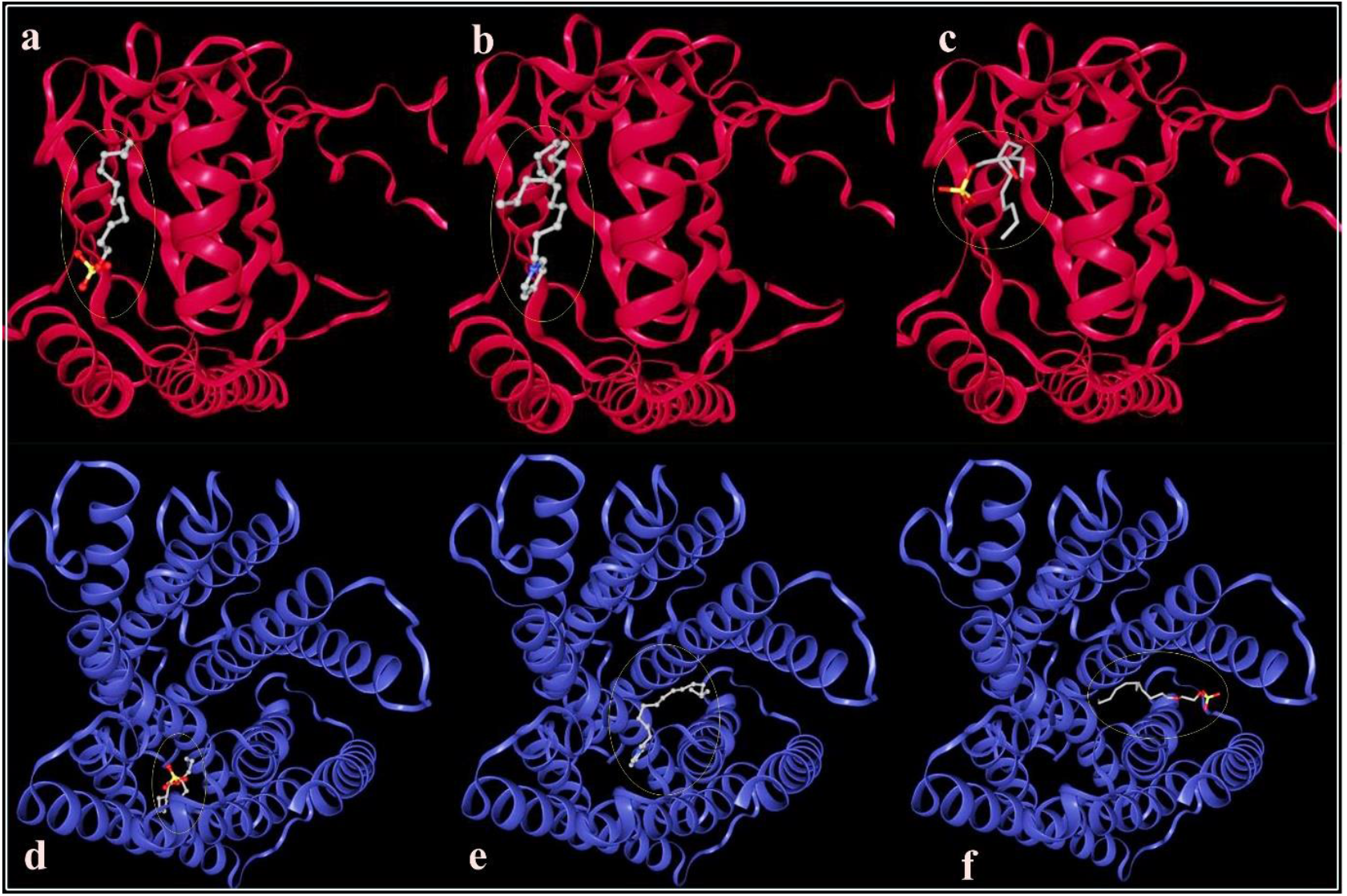
3D structure of docking between **(a):** Cytochrome c oxidase subunit 2 of *Cyprinus carpio* and SDS, **(b):** Cytochrome c oxidase subunit 2 of *Cyprinus carpio* and CPC, **(c):** Cytochrome c oxidase subunit 2 of *Cyprinus carpio* and SLES, **(d):** Cytochrome c oxidase subunit 1 of *Tubifex tubifex* and SDS **(e):** Cytochrome c oxidase subunit 1 of *Tubifex tubifex* and CPC, **(f):** Cytochrome c oxidase subunit 1 of *Tubifex tubifex* and SLES. The ball and stick structure represent the ligand.

Vina score is a quantitative scoring function that determines the affinity, or efficiency, of protein-ligand interactions (Liu and Wang 2015). Low Vina scores indicate high binding affinity between protein and ligand (Wójcikowski *et al.* 2017). The vina scores of surfactant protein complex is displayed in **Table 2**. Surfactant protein interaction profile analysis are mainly focused on hydrogen bonds and hydrophobic interaction and to some extent salt bridge. Hydrogen bonds greatly influence the value of binding affinity. The strength of hydrogen bonds can be subdivided dependent on the distance between the interactions. Hydrogen bonds with a distance of 2.2–2.5 Å are considered strong, 2.5–3.2 Å are moderate, and 3.2–4.0 Å are weak (Jeffrey 1997, Khayrani *et al.* 2021). In terms of the interaction between the protein and its ligand, hydrophobic interactions are the most dominant (Khayrani *et al.* 2021). Referencing to literature, several standard values of distance cutoffs to consider the formation of hydrophobic interactions have been found and the optimum range is in between 3.3 – 3.8 Å (Janiak 2000). While other researchers have suggested a relatively higher ranges (Burley and Petsko 1985, Peikert *et al.* 2014, Piovesan *et al.* 2016). The interactions and active site residues responsible for bindings between surfactants and target protein Cytochrome c oxidase subunit 2 of *Cyprinus carpio* and Cytochrome c oxidase subunit 1 of *Tubifex tubifex* are presented in **Fig 8** and **9** respectively.

**Table 2:**
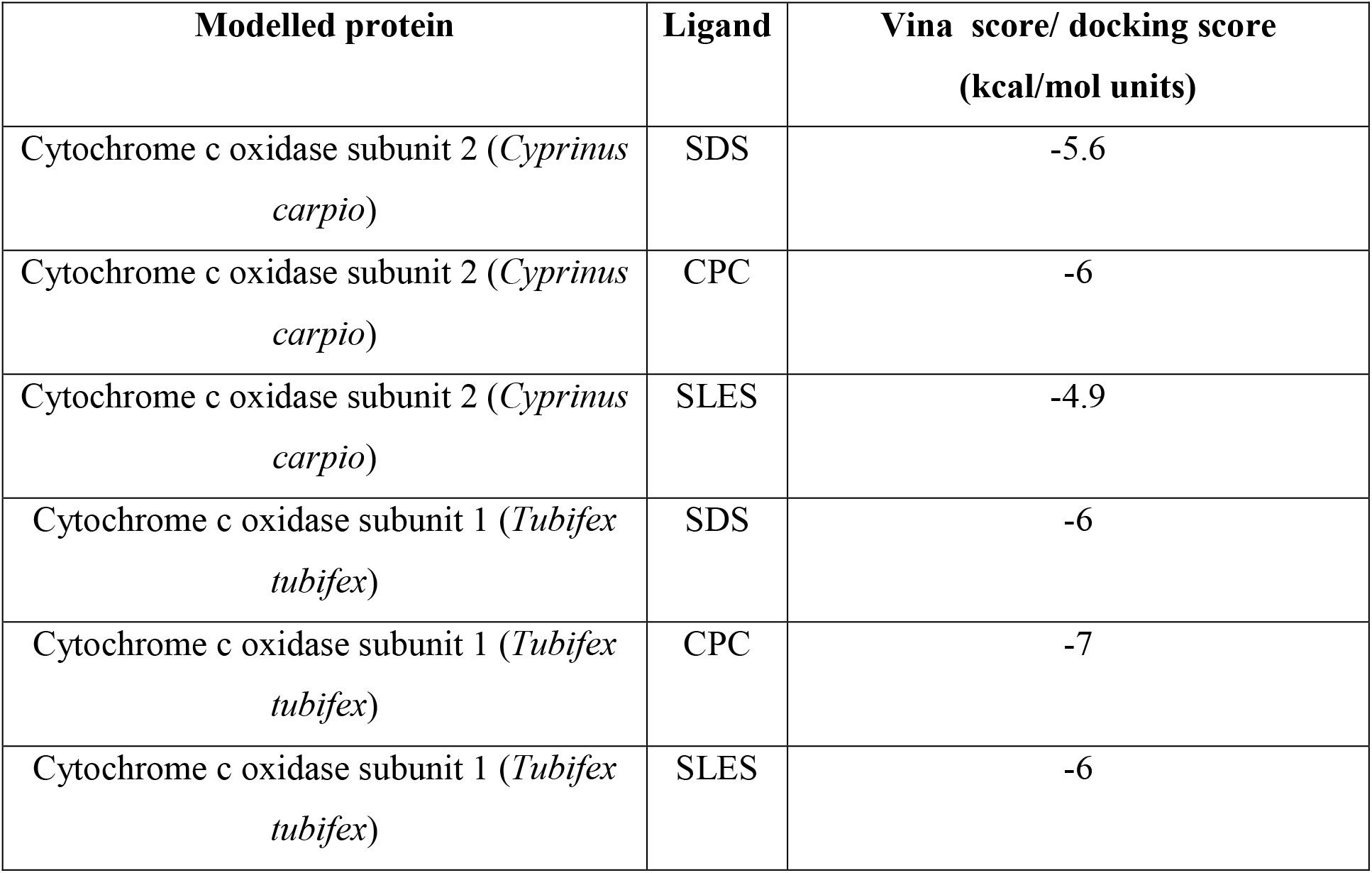
Vina score between modelled protein and surfactants

| Modelled protein | Ligand | Vina score/ docking score<br>(kcal/mol units) |
| --- | --- | --- |
| Cytochrome c oxidase subunit 2 (*Cyprinus carpio*) | SDS | -5.6 |
| Cytochrome c oxidase subunit 2 (*Cyprinus carpio*) | CPC | -6 |
| Cytochrome c oxidase subunit 2 (*Cyprinus carpio*) | SLES | -4.9 |
| Cytochrome c oxidase subunit 1 (*Tubifex tubifex*) | SDS | -6 |
| Cytochrome c oxidase subunit 1 (*Tubifex tubifex*) | CPC | -7 |
| Cytochrome c oxidase subunit 1 (*Tubifex tubifex*) | SLES | -6 |

**Fig 8:**
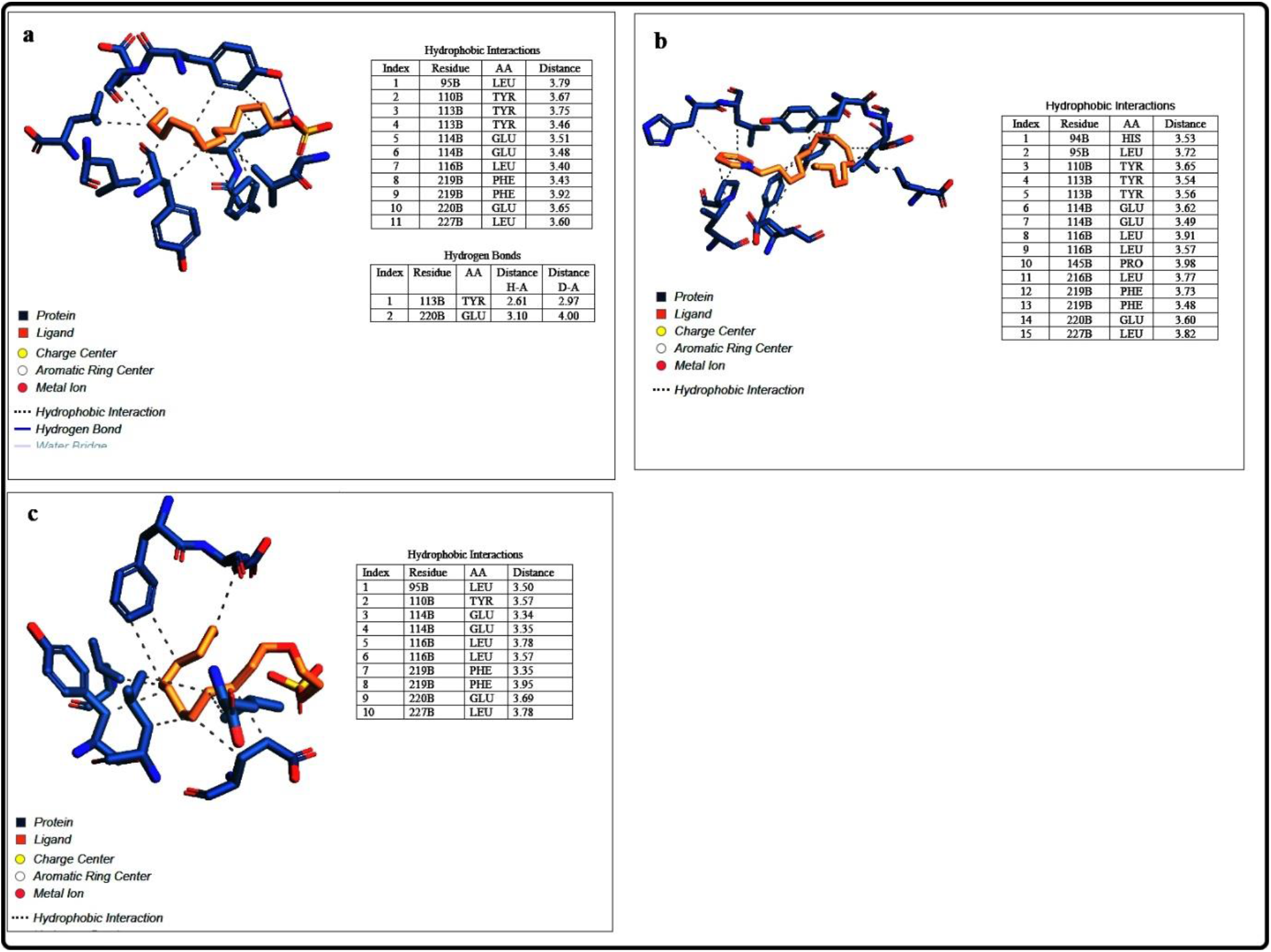
Types of interaction between **(a):**Cytochrome c oxidase subunit 2 of *Cyprinus carpio* and SDS, **(b):** Cytochrome c oxidase subunit 2 of *Cyprinus carpio* and CPC, **(c):** Cytochrome c oxidase subunit 2 of *Cyprinus carpio* and SLES.

**Fig 9:**
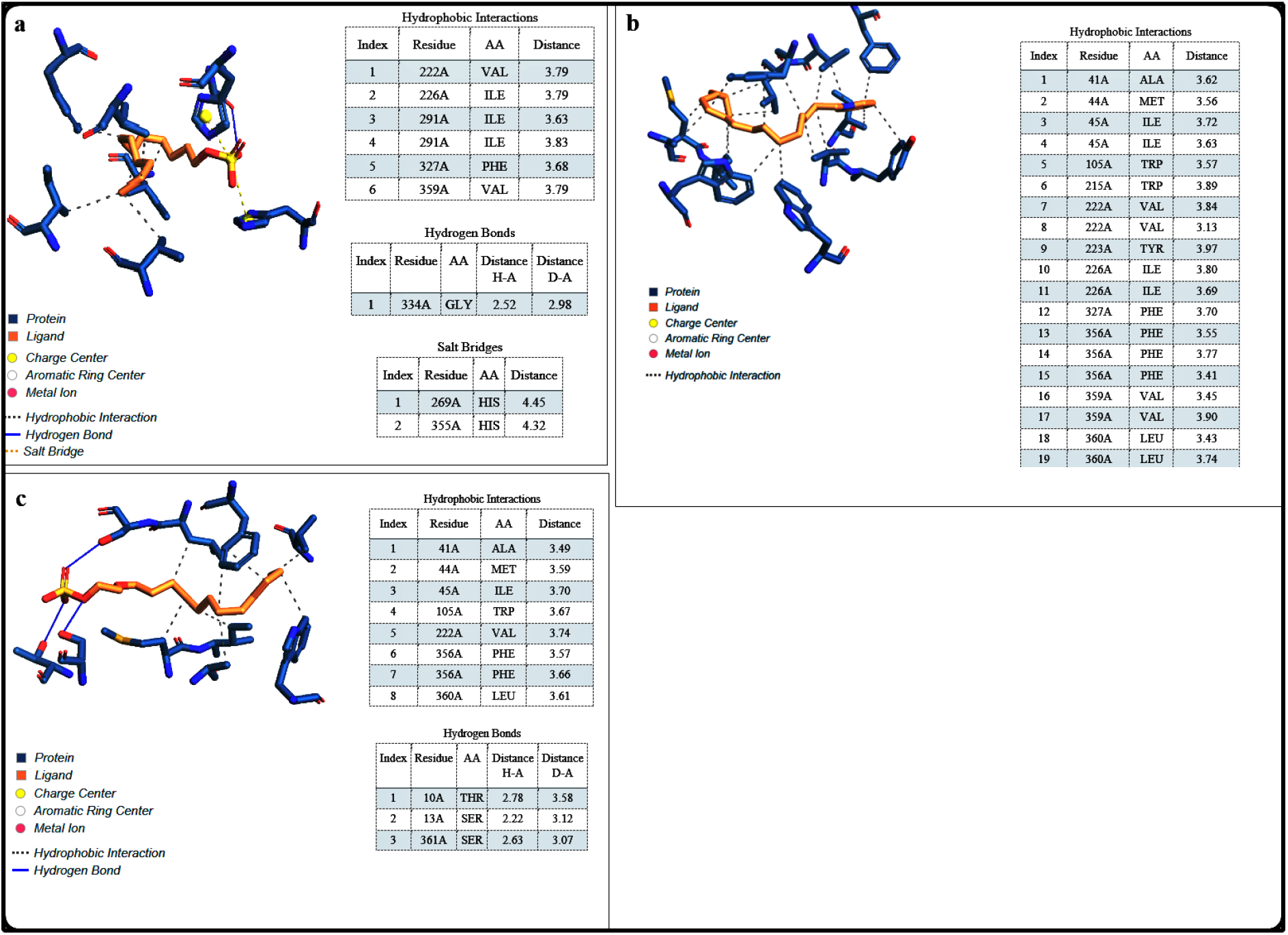
Types of interaction between **(a):** Cytochrome c oxidase subunit 1 of *Tubifex tubifex* and SDS, **(b):** Cytochrome c oxidase subunit 1 of *Tubifex tubifex* and CPC, **(d):** Cytochrome c oxidase subunit 1 of *Tubifex tubifex* and SLES.

In the case of Cytochrome c oxidase subunit 2 of *Cyprinus carpio* SDS demonstrated the vina score of −6 kcal/mol associated with hydrophobic interaction with the residues of Leu 95, Tyr 110, Tyr 113, Glu 114, Leu 116, Phe 219, Glu 220, Leu 227, and hydrogen bonding with residues Tyr 113 and Glu 220 as well as salt bridge interaction with residues His 269 and 355. CPC demonstrated the vina score of −7 kcal/mol with hydrophobic interaction with the residues of His 94, Leu 95, Tyr 110, Tyr 113, Glu 114, Leu 116, Pro 145, Leu 216, Phe 219, Glu 220, Leu 227. SLES demonstrated the vina score of −6 kcal/mol with hydrophobic interaction with the residues of Leu 95, 116, 227, Tyr 110, Glu 114, Phe 219 and Glu 220. Moreover in case of Cytochrome c oxidase subunit 1 of *Tubifex tubifex* SDS demonstrated the vina score of −6 kcal/mol associated with hydrophobic interaction with the residues of Val 222, 359 Ile 226, 291 and Phe 327 and hydrogen bonding with residues Gly 334. CPC demonstrated the vina score of −7 kcal/mol with hydrophobic interaction with the residues of Ala 41, Met 44, Ile 45, 226, Trp 105, 215, Val 222, 359, Tyr 223, Phe 327, 356 and Leu 360.

Moreover, SLES demonstrated the vina score of −6 kcal/mol with hydrophobic interaction with the residues of Ala 41, Met 44, Ile 45, Trp 105, Val 222, Phe 356, Leu 360, and hydrogen bonding with residues Thr 10, Ser 13 and 361. However, based on the vina score, it is fairly obvious that the surfactant CPC interacted the best with Cytochrome c oxidase subunit 2 of *Cyprinus carpio* and Cytochrome c oxidase subunit 1 of *Tubifex tubifex*. SDS and SLES also interact with the modelled proteins, albeit to a lower extent than CPC. Thus these results are in good concordance with the outcome of several researchers who reported that cationic surfactants are more toxic than anionic surfactants in aquatic organisms (Jardak *et al.* 2016). This might be due to this increased binding affinity of cationic surfactants to Cytochrome c oxidase than anionic surfactants. In our study in terms of the interaction between the protein and surfactants, maximum number and occurrence of hydrophobic interactions with the active site residues were observed rather than hydrogen bonds and salt bridges and the maximum existence of this hydrophobic interactions within the catalytic sites may indicate that the surfactants can proceed with inhibitory activity (Khayrani *et al.* 2021).

### 7. Molecular dynamics (MD) studies

To explore the structural and dynamic changes, we identified the residues regions of Cytochrome c oxidase (*Tubifex tubifex*) and Cytochrome c oxidase (*Cyprinus carpio*) either in unbounded or bounded with ligands form by analyzing the structural mobility based on the RMSF of the backbone atoms with respect to the initial structure. The conformational consistency of the docked poses is expressed by the RMSF values. The higher RMSF most likely are loop regions with more conformational flexibility, whereas the lower value reflects the proteins’ restricted movement (Shah *et al.* 2021). As per our findings, the maximum RMSF value for both enzymes: Cytochrome c oxidase (*Cyprinus carpio*) and Cytochrome c oxidase (*Tubifex tubifex*) in unbounded cases are: 7.8 Å and 6.1 Å respectively **[Figure 10(a,e)].** On the other hand, we can observe that the RMSF values obtained in the bounded structure are low compared to the unbounded form **[Figure 10 (b, c, d, f, g, h)].**

**Fig 10:**
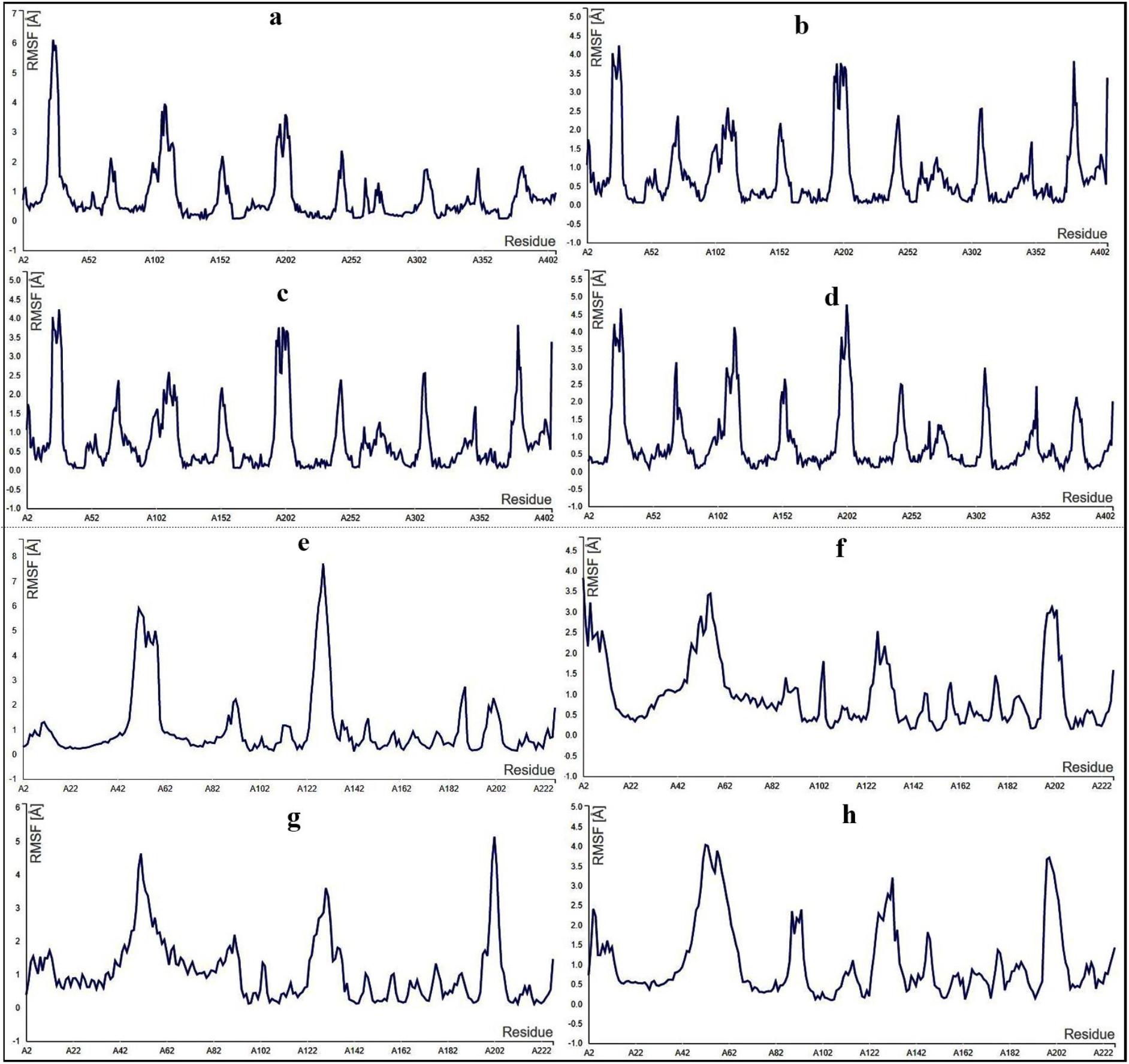
Molecular dynamics simulation showing the RMSF of **(a)** an unbounded Cytochrome c oxidase (*Tubifex tubifex*) **(b)** Cytochrome c oxidase (*Tubifex tubifex*) bound to SDS **(c)** Cytochrome c oxidase (*Tubifex tubifex*) bound to SLES **(d)**Cytochrome c oxidase (*Tubifex tubifex*) bound to CPC **(e)**unbounded Cytochrome c oxidase of *Cyprinus carpio* **(f)**Cytochrome c oxidase (*Cyprinus carpio*) bound to SDS **(g)**Cytochrome c oxidase (*Cyprinus carpio*) bound to SLES **(h)**Cytochrome c oxidase (*Cyprinus carpio*) bound to CPC.

Cytochrome c oxidase (*Cyprinus carpio*) in unbounded form exhibited several fluctuations with the presence of high amplitudes at residues: 51 (RMSF≈6 Å) and 129 (RMSF≈7.8Å). In addition, the RMSF values of Cytochrome c oxidase (*Cyprinus carpio*) in SDS bounded form indicated a slight fluctuation in regions of the residues: 5 (RMSF≈3.4 Å), 56 (RMSF≈3.5 Å) and 200 (RMSF≈3.2 Å) **[Fig10(f)]**.Also, the RMSF values of Cytochrome c oxidase (*Cyprinus carpio*) in SLES bounded are low in the residues: 51 (RMSF≈4.6 Å) and 130 (RMSF≈3.6 Å) but exception in residue: 202 (RMSF≈5.2 Å) which showed steep RMSF fluctuations in this residue **[Fig10(g)]**. Finally, the RMSF values of Cytochrome c oxidase (*Cyprinus carpio*) in CPC bounded form are low in the following residues: 52 (RMSF≈4.1 Å), 57 (RMSF≈3.9 Å), 132 (RMSF≈3.3 Å) and 200 (RMSF≈3.7 Å) **[Fig10(h)]**.

In the case of *Tubifex tubifex*, several fluctuations are present with the persistence of high amplitudes in different residues of Cytochrome c oxidase [25 (RMSF≈6.1 Å and 111 (RMSF≈3.8 Å] in unbounded form **[Fig 10(a)]**. Residue wise comparison exhibited that the RMSF values of the same residues 25 and 111 is much lower in SDS bounded [25 (RMSF≈3.3 Å, 111 (RMSF≈2.3 Å)], SLES bounded [(25 (RMSF≈3.4 Å, 111 (RMSF≈2.3 Å)] and CPC bounded [25 (RMSF≈3.7 Å, 111 (RMSF≈2.6 Å)] form **[Fig. 10 (b, c and d)]**.

Thus our results indicate that the binding of the residues of protein to the ligands affects the local structure and conformation of the protein.

## Conclusion

The result suggests that the commonly used surfactants Sodium dodecyl sulphate, Cetylpyridinium chloride and Sodiun laureth sulphate might impart detrimental effect on the structure and dynamics of Cytochrome c oxidase in fish and fish food organism by binding with it in varying degrees. Due to this preliminary encouraging result based on Insilico assessment, further studies are required in the field trials for validation of our findings.

## Ethical approval

This study does not include animal experiments by the authors that require the ethics committee’s permission.

## Acknowledgment

The authors are thankful to the Department of Zoology, The University of Burdwan for giving all sorts of technical and computational facilities to conduct this research.

